# Scanning mutagenesis of RNA-binding protein ProQ reveals a quality control role for the Lon protease

**DOI:** 10.1101/2021.07.12.452043

**Authors:** Youssef El Mouali, Falk Ponath, Vinzent Scharrer, Nicolas Wenner, Jay C. D. Hinton, Jörg Vogel

## Abstract

The FinO-domain protein ProQ belongs to a widespread family of RNA-binding proteins (RBPs) involved in gene regulation in bacterial chromosomes and mobile elements. Whilst the cellular RNA targets of ProQ have been established in diverse bacteria, the functionally crucial ProQ residues remain to be identified under physiological conditions. Following our discovery that ProQ deficiency alleviates growth suppression of *Salmonella* with succinate as the sole carbon source, an experimental evolution approach was devised to exploit this phenotype. By coupling mutational scanning with loss-of-function selection, we identified multiple ProQ residues in both the N-terminal FinO domain and the variable C-terminal region required for ProQ activity. Two C-terminal mutations abrogated ProQ function and mildly impaired binding of a model RNA target. By contrast, several mutations in the FinO domain rendered ProQ both functionally inactive and unable to interact with target RNA *in vivo*. Alteration of the FinO domain stimulated the rapid turnover of ProQ by Lon-mediated proteolysis, suggesting a quality control mechanism that prevents the accumulation of non-functional ProQ molecules. We extend this observation to Hfq, the other major sRNA chaperone of enteric bacteria. The Hfq Y55A mutant protein, defective in RNA-binding and oligomerization, proved to be labile and susceptible to degradation by Lon. Taken together, our findings connect the major AAA+ family protease Lon with RNA-dependent quality control of Hfq and ProQ, the two major sRNA chaperones of Gram-negative bacteria.

**SIGNIFICANCE:** Proteins that interact with RNA play a vital role in controlling key functions in pathogenic bacteria. RNA-binding proteins regulate how, when and where bacteria feed, swim or interact with a host, and it is critical that we understand how RNAs associate with these proteins. ProQ is one of the three major RNA-binding proteins (RBPs) in Gram-negative bacteria. In this study, we mapped the amino acid residues of ProQ that are essential for function. We successfully identified residue substitutions that rendered the ProQ RBP both non-functional and unable to interact with RNA. Our findings raise the possibility that the Lon protease mediates a quality control mechanism of ProQ that targets this RBP in the absence of RNA. A posttranslational quality control mechanism of this type could prevent the accumulation of nonfunctional RBPs in the bacterial cytoplasm.

## INTRODUCTION

Globally acting RNA-binding protein (RBPs) that work in concert with small regulatory RNAs (sRNAs) play crucial roles in post-transcriptional control in both eukaryotes and prokaryotes (1). Three RBPs are highly conserved in Gram-negative bacteria (2): the translational repressor CsrA (a.k.a. RsmA), which itself is regulated by decoy sRNAs (3), and the RNA chaperones Hfq and ProQ, both of which regulate mRNAs via base pairing sRNAs (3–7). The *in vivo* targetomes of these central RBPs have been mapped extensively in several bacteria (8, 9, 18–21, 10–17), revealing that each interacts with hundreds of different transcripts from diverse cellular pathways. However, whilst their regulatory roles and their associated sRNAs become better understood, we still know little about the biogenesis and turnover of these central RPBs, and the mechanisms that ensure that they do not accumulate intracellularly as non-functional RBPs. In this study, we propose a quality control mechanism that involves the degradation of ProQ when it fails to associate with RNA.

ProQ is the least understood of the three global RBPs. In *Salmonella enterica* serovar Typhimurium (henceforth, *Salmonella*), ProQ is an abundant 25 kDa protein with an N-terminal domain (NTD, residues 1-119) and a C-terminal domain (CTD, 176-228 residues) connected by a disordered ∼50-aa linker region (9, 22). The targets of ProQ include ∼50 sRNAs that tend to be more structured than Hfq-bound sRNAs (9, 15), and several hundred mRNAs, which are primarily recognized at their 3’ end (15); many of these interactions are conserved between *Salmonella* and *Escherichia coli* (14, 15). Target recognition is thought to be determined by the NTD, a region that contains a FinO domain (PFAM04352) and was shown to mediate RNA binding of other investigated members of the family of ProQ/FinO-like proteins (11, 12, 23–25). In addition, the NTD was sensitive to mutations in a bacterial three-hybrid (B3H) assay designed to score ProQ binding to selected bait RNAs (26). By contrast, the role of the CTD is unclear. A truncated ProQ protein lacking the variable C-terminal region showed weaker binding to some targets *in vitro* as compared to full-length ProQ (27), but the importance of the CTD for *in vivo* activity of ProQ is yet to be demonstrated.

Forward genetics, using saturation mutagenesis coupled with phenotypic screening, is a powerful tool to assess the importance of individual regions and amino acid residues for the *in vivo* function of a protein of interest (28). However, despite the established global activity of ProQ as an RBP, the known phenotypes of *Salmonella* Δ*proQ* strains (e.g., impaired motility and reduced invasion of eukaryotic cells (29)) were insufficiently robust for *in vivo* screens under physiological conditions. Inspired by the reported growth suppression of *Salmonella* with succinate as sole carbon source (30), we discovered that deletion of the *proQ* gene generated a gain-of-function phenotype that permitted rapid growth on minimal media containing succinate. Using this strong phenotype for a saturation mutagenesis screen, we have mapped crucial residues for ProQ function *in vivo*. We establish a link between RNA binding ability and protein stability, suggesting a quality control mechanism in which non-functional ProQ is rapidly removed from the cell by the major ATP-dependent protease Lon.

## RESULTS

### ProQ suppresses growth in succinate-containing media

Succinate has been identified as an important carbon source for *Salmonella* during colonization of the murine gastrointestinal tract (31), and plays an important role during macrophage infection (32, 33). Wild-type *S*. Typhimurium displays an extended lag phase and a slow doubling time when grown in media with succinate as the sole carbon source (30). Because deletion of the *S*. Typhimurium *proQ* gene is known to modulate expression of hundreds of mRNAs and sRNAs (9), we investigated whether ProQ modulated the succinate-dependent *in vitro* growth phenotype. We discovered that genetic inactivation of *proQ* increases the growth rate of *Salmonella* when succinate is the sole carbon source, either on agar plates or under microaerobic conditions in liquid media (Fig. 1A-B).

**Figure 1.**
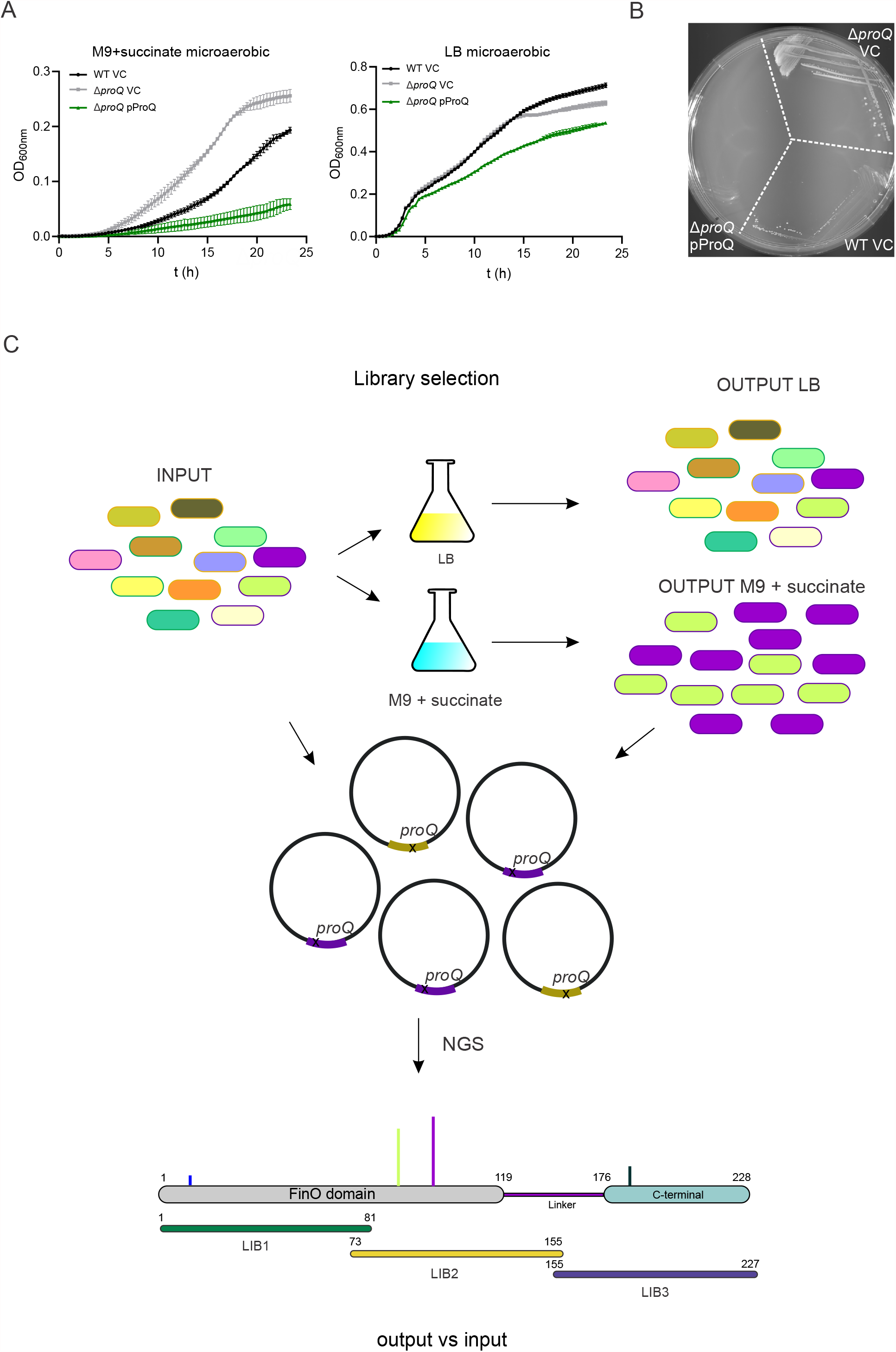
ProQ represses succinate utilization in *Salmonella*. **A**. Growth of *Salmonella* WT, Δ*proQ* and Δ*proQ* pProQ in a 96-well plate Tecan. Strains were inoculated to an OD_600 nm_ 0.01 and grown in a 96-well plate in M9 40mM succinate (or LB) at 37°C for 24h without shaking. VC: presence of the empty pZE12vector (pJV300) **B**. Growth of *Salmonella enterica* SL1344 WT and Δ*proQ* derivative strain in solid agar media M9 with 40 mM succinate as sole carbon source. Single colony was streaked and plates incubated overnight at 37 °C. **C**. Deep mutational scanning schematic workflow. The input represents a library of mutants expressed from pZE12-ProQ (pProQ) within a Δ*proQ* strain. The libraries were selected in parallel in non-selective media (LB) and selective media (M9 40mM Na-succinate). The cultures were incubated without shacking at 37°C for 24h. Form the resulting cultures, plasmid content was extracted and used as template for library preparation and Mi-seq 2×300bp sequencing. In the bottom, schematic representation of ProQ protein organization and coordinates of ProQ mutant libraries. LIB1 1-81 aa, LIB2 73-155 aa and LIB3 155-227 aa.

This gain-of-function phenotype is shown in the well-characterized Δ*proQ* strain of *S*. Typhimurium SL1344 (9, 23, 29), and we confirmed that plasmid complementation fully restored the succinate-dependent growth suppression by ProQ (Fig. 1A-B). The molecular basis of this conditional growth permissiveness is unknown, but our preliminary experiments point towards a derepression of the TCA cycle in the absence of ProQ (Fig. S1). The remarkable robustness of this succinate-dependent growth phenotype provided the first unequivocal readout of functionality of *Salmonella* ProQ *in vivo*.

### Deep mutational scanning of ProQ in vivo

To obtain a functional map of ProQ residues *in vivo*, we performed saturation mutagenesis of the *Salmonella proQ* open reading frame (ORF). To this end, we generated libraries of ProQ expression plasmids (pProQ, *proQ* expressed under its native promoter) with random mutations in the coding sequence (CDS) of *proQ* that were introduced into the Δ*proQ* strain. The plasmids that allowed microaerobic growth in M9-succinate media were sequenced to identify amino acid substitutions in the ProQ protein. Importantly, wild-type *Salmonella*, the Δ*proQ* strain alone or the latter complemented with *proQ* on a plasmid all displayed similar fitness when grown in LB media (Fig. 1A), which provided a neutral growth condition for constructing and maintaining these mutant libraries.

Mutant variants of *proQ* were generated by error prone PCR, cloned and expanded in *E. coli* TopF cells to ultimately be transformed within the Δ*proQ* strain (Fig. S2). To overcome the limited read length of Illumina sequencing (300 bp pair-end sequencing), we generated three mutant libraries that focused on different regions of the 228-aa ProQ protein: LIB1, LIB2, and LIB3, covering amino acids 1-81, 73-155, and 155-227, respectively (Fig. 1C). The mutants for each of the libraries were combined individually to obtain ‘input’ pools for LIB1, LIB2, and LIB3, respectively. We then screened a total of 30,000 individual colonies (containing ∼5,000 variants, see methods) for each of the ‘input’ pools for growth (i.e., loss of ProQ function) in liquid M9-succinate media for 24 h without aeration (Fig. 1C). In parallel, the ‘input’ pools were grown in LB to determine whether ProQ variants accumulated due to genetic drift, independent of the selection for ProQ functionality. We will refer to these pools from growth in LB or M9-succinate media as ‘output LB’ or ‘output M9’, respectively (Fig. 1C).

Sequencing the input and output pools, we detected thousands of variants, including 93 mutants that carried stop mutations. Only the single non-synonymous substitutions were considered in the biocomputational analysis: 1,056 mutants for LIB1; 1,164 for LIB2; and 1,032 for LIB3. Of note, single substitutions for all individual residues of ProQ were detected in both the input (Supplementary Table S2) and output libraries. For each of the three different libraries, a prominent enrichment (Z-score ≥ 30, enrich score ≥ 1.4) of amino acid substitutions in the output versus the input was only observed after selection in M9-succinate media (Fig. 2A-F). Thus, these enriched residues represented candidate mutants of ProQ protein with impaired function *in vivo*.

**Figure 2.**
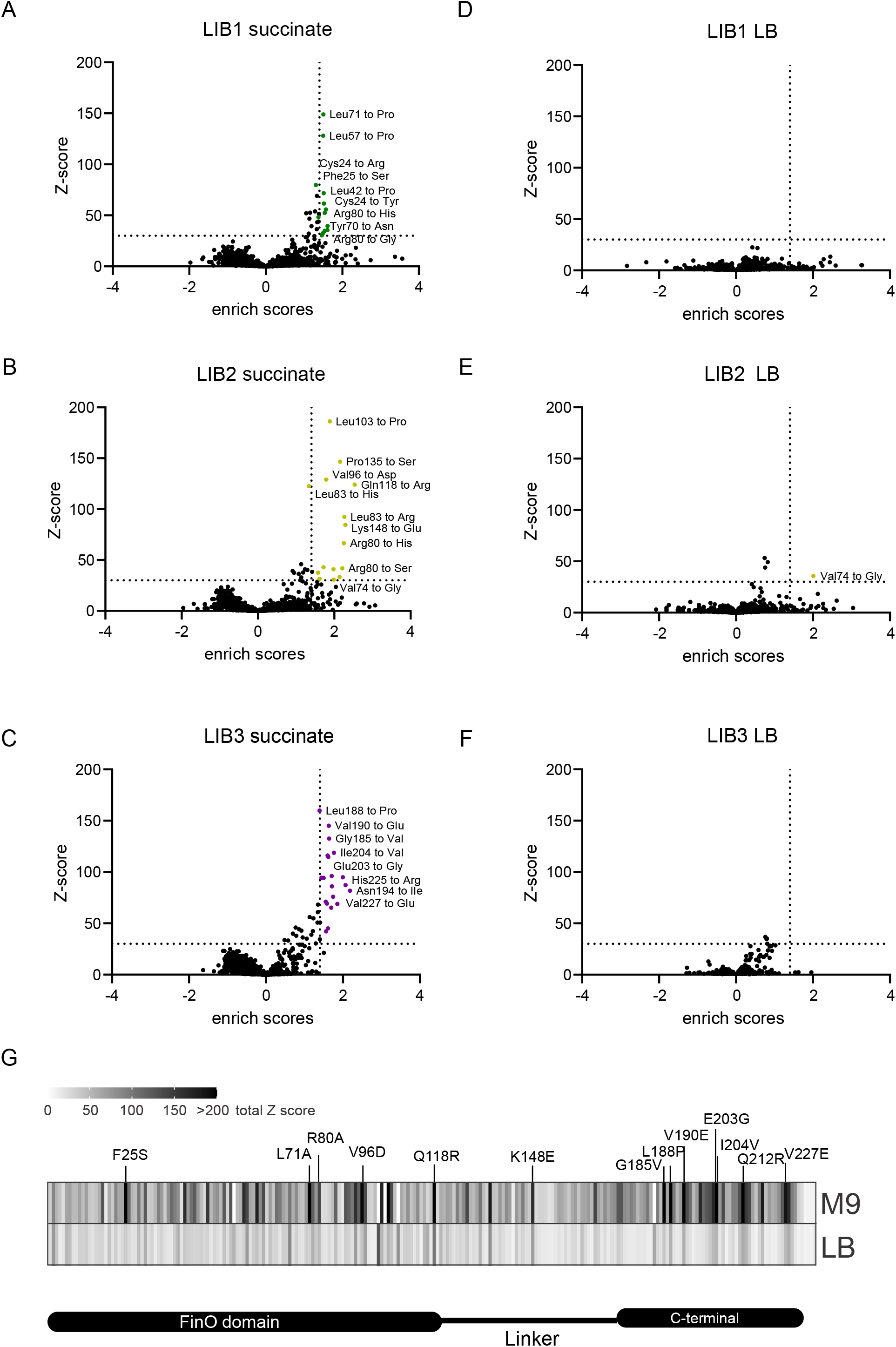
ProQ variants enrichment. **A**. Amino acid substitutions enriched in the output in M9 succinate (selective media) when compared to the input for LIB1 (1-81 aa). In green highlighted significantly enriched residues substitutions. **B**. Amino acid substitutions enriched in the output in M9 succinate (selective media) when compared to the input for LIB2 (73-155 aa). In yellow highlighted significantly enriched residues substitutions. **C**. Amino acid substitutions enriched in the output in M9 succinate (selective media) when compared to the input for LIB3 (155-227 aa). In purple highlighted significantly enriched residues substitutions. **D**. Amino acid substitutions enriched in the output in LB (non-selective media) when compared to the input for LIB1 (1-81 aa). **E**. Amino acid substitutions enriched in the output in LB (non-selective media) when compared to the input for LIB2 (73-155 aa). In yellow highlighted significantly enriched residues substitutions. **F**. Amino acid substitutions enriched in the output in LB (non-selective media) when compared to the input for LIB3 (155-227 aa). **G**. Heatmap summarizing panels A-F. For each residue, the Z-score of all detected substitutions was summed and represented. Higher values indicate strong enrichment as loss-of-function mutant in the given position.

### Both the N-terminus and C-terminus of ProQ are essential for in vivo function

LIB1 and LIB2 collectively cover the N-terminal FinO domain of ProQ, and all the 27 top-enriched (Z-score ≥ 30, enrich score ≥ 1.4) mutations from the output M9 pool fall within this domain, except ProQ_P135S_ and ProQ_K148E_ in the linker region (Fig. 2G). Importantly, although the selected residues appeared to be scattered over the FinO domain in terms of primary sequence, they spatially clustered in the tertiary structure and overlapped with the proposed RNA interaction face of ProQ (see further below). Intriguingly, the third library (LIB3) also predicted numerous residues important for ProQ function; these clustered in a 40-aa region that overlaps with a predicted C-terminal Tudor domain in ProQ (22), while none were selected in the upstream linker region (Fig. 2G). Overall, the distribution of all these enriched mutations supports the current coarse-grained structural model of ProQ as a protein with two distinguishable functional domains, separated by a variable linker region (22).

To validate our screening results, we selected and individually introduced 13 of the highly enriched mutants in a ProQ expression plasmid. All mutant proteins carried a C-terminal 3×FLAG epitope, added to ensure unbiased western blot detection and to permit immunoprecipitation with a generic α-FLAG antibody. Following the introduction of these plasmids into *Salmonella* Δ*proQ*, we evaluated growth in M9-succinate, expecting they would phenocopy lack of ProQ (Fig. 3A-B). We found that four (ProQ_F25S_, ProQ_L71A_, ProQ_R80A_, and ProQ_V96D_) of the five tested mutations in the FinO domain did permit growth, whereas the ProQ_Q118R_ mutation still repressed growth under this non-permissive condition (Fig. 3A). Mutation ProQ_K148E_ in the linker region did not affect ProQ-mediated growth repression (Fig. 3A).

**Figure 3.**
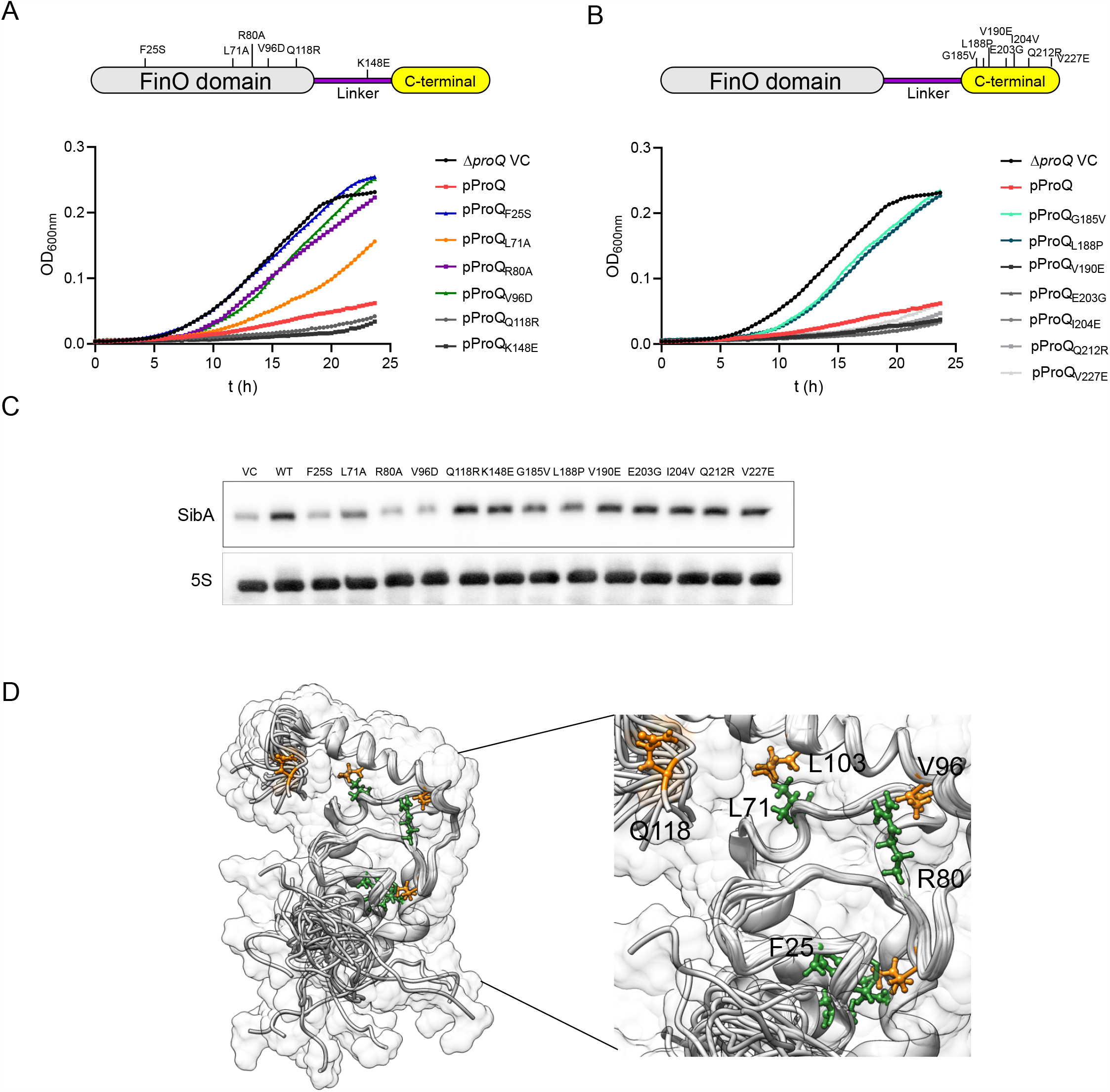
Identification of ProQ non-functional variants. **A**. Upper panel, schematic representation of ProQ protein domains. Site directed mutants generated in ProQ are indicated. ProQ_F25S_, ProQ_L71A_, ProQ_R80A_, ProQ_V96D_, ProQ_Q118R_ and ProQ_K148E_. Bottom panel, growth curve of Δ*proQ* strain carrying either vector control (VC), pProQ WT protein or generated ProQ mutant variants indicated in the upper panel. **B**. Upper panel, schematic representation of ProQ protein domains. Site directed mutants generated in ProQ are indicated. ProQ_G185V_, ProQ_L188P_, ProQ_V190E_, ProQ_E203G_, ProQ_I204E_, ProQ_Q212R_ and ProQ_V227E_. Bottom panel, growth curve of Δ*proQ* strain carrying either VC or the generated ProQ mutant variants indicated in the upper panel. Strains were inoculated to an OD_600 nm_ 0.01 and grown in a 96-well plate in M9 40mM succinate at 37°C for 24h without shaking. **C**. Northern blot detection of SibA expression in Δ*proQ* strain carrying pProQ WT or generated mutant variants as in A. and B. 5S RNAwas detected as loading control. Total RNA samples were obtained from cultures grown in LB to OD_600nm_ 2.0. **D**. Representation of ProQ NTD (FinO domain) (PDB ID: 5nb9). Residues enriched deep mutational scanning are highlighted in green (LIB1) or orange (LIB2). On the right, highlighted cluster within ProQ structure of residues whose substitution render protein unstable, ProQ_F25S_, ProQ_R80A_ ProQ_L71A,_ ProQ_V96D._

In regard to the C-terminal domain, two (ProQ_G185V_, ProQ_L188P_) of the seven tested point mutations passed this independent validation step, failing to repress growth in succinate (Fig. 3B). Of note, whilst all mutants in LIB3 were assumed to possess a wild-type FinO domain, we cannot rule out mutations in the FinO domain because the screen only involved sequencing of the mutagenized region of LIB3 (155-227). This possibility of additional mutations in other regions might explain the lower validation rate of C-terminal point mutations. Nevertheless, our mutagenesis approach identified non-functional variants of ProQ via a single amino acid substitution in either the FinO domain or the C-terminal domain.

### Effects of ProQ mutations on target RNA levels

Assuming that ProQ primarily works as an RBP, we next tested whether the confirmed six mutations affected the ability of ProQ to interact with cellular RNA targets. We used the SibA sRNA as a proxy, which is a top ligand of ProQ *in vivo* (9, 15, 23). Importantly, SibA must be strongly associated with ProQ to be stable in the cell (9, 29), providing an easy readout for intact RNA recognition. Northern blot analysis revealed lower levels of SibA in the presence of mutants ProQ_F25S_, ProQ_L71A_, ProQ_R80A_ and ProQ_V96D_, compared with the Δ*proQ* strain expressing wild-type ProQ; in fact, these mutants showed SibA levels close to the Δ*proQ* strain carrying the empty control vector (Fig. 3C). In other words, all four mutations that were unable to repress growth on succinate did not sustain normal SibA levels. Intriguingly, the ProQ_F25S_, ProQ_L71A_ and ProQ_R80A_ substitutions were also predicted to be crucial for RNA binding in the *E. coli* 3HB screen (26), and these residues are in close vicinity to one another on the assumed RNA binding face of ProQ (Fig. 3D).

By contrast, the two confirmed C-terminal mutants, ProQ_G185V_ and ProQ_L188P_, displayed minor differences in SibA RNA levels when compared to the strain expressing wild-type ProQ, despite the loss-of-function in growth repression on succinate. These differences suggest that the FinO domain and the C-terminal domain seem to play different roles in control of the succinate phenotype, with the N-terminal mutations being more likely to generally impair ProQ association with cellular target transcripts.

### RNA-binding deficient ProQ proteins are unstable in vivo

While the lower SibA levels might simply be explained by loss-of-function of key residues for RNA binding, we double-checked another possible cause, i.e., reduced protein levels of these ProQ variants. After probing the expression levels of 13 of the mutant proteins on a western blot (Fig. 4A), we observed dramatically lower levels of the exact same four FinO domain mutants (ProQ_F25S_, ProQ_L71A_, ProQ_R80A_ and ProQ_V96D_) accompanied by reduced SibA levels and confirmation of the loss of growth suppression on succinate. In contrast, all other seven mutants showed at least wild-type protein levels, with the two C-terminal mutants ProQ_G185V_ and ProQ_L188P_ accumulating to even higher levels (Fig. 4A).

**Figure 4.**
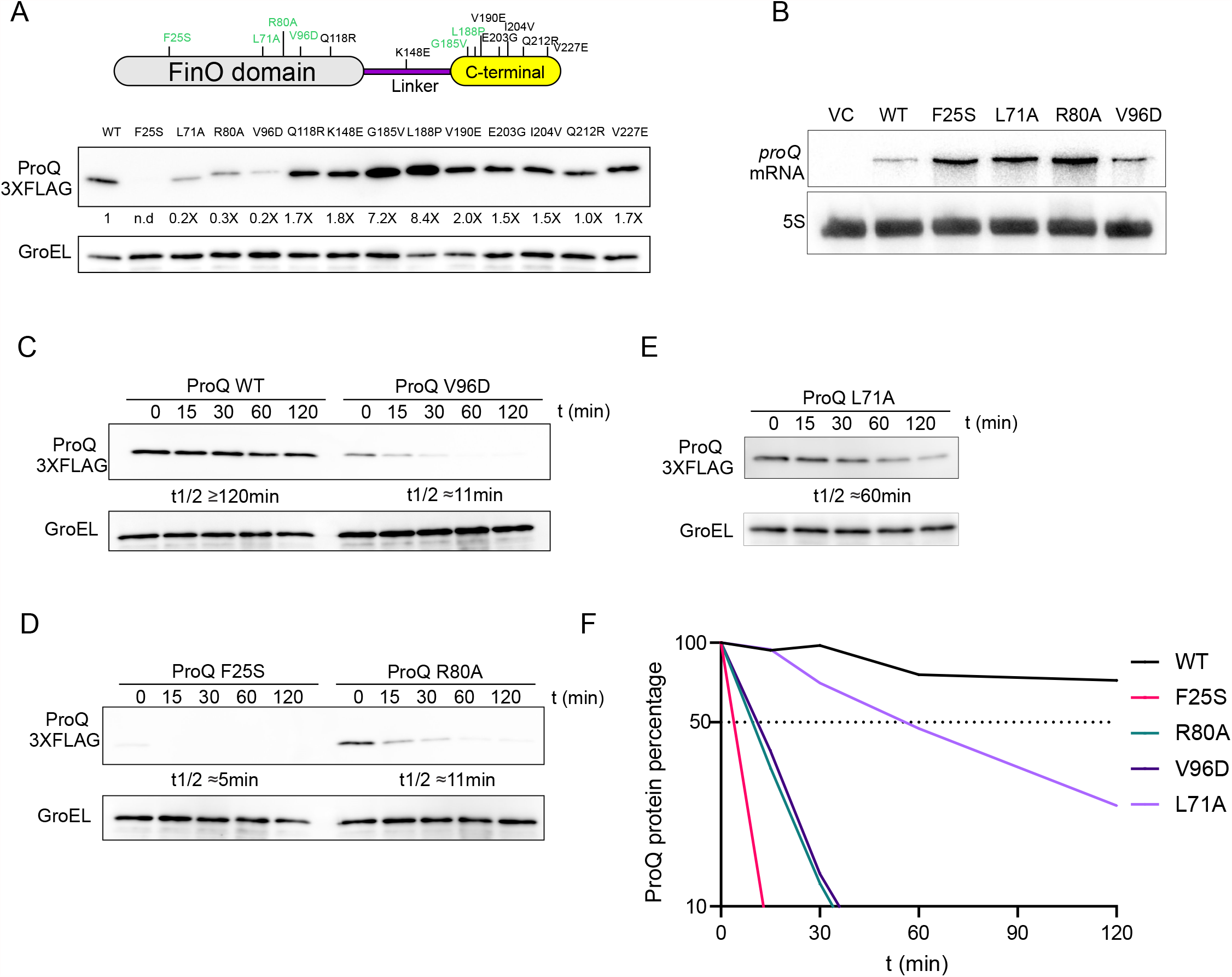
Mutations in the FinO domain render ProQ unstable. **A**. Upper panel, schematic representation of ProQ protein domains. Site directed mutants generated in ProQ are indicated. In green, ProQ variants that are not functional. ProQ_F25S_, ProQ_L71A_, ProQ_R80A_, ProQ_V96D_, ProQ_G185V_ and ProQ_L188P_. Bottom panel, western blot of Δ*proQ* strain carrying either pProQ-3×FLAG WT protein or generated ProQ mutant variants indicated in the upper panel. GroEL was immunodetected as loading control. **B**. Northern blot detection of *proQ* mRNA expression of pProQ WT, ProQ_F25S_, ProQ_L71A_, ProQ_R80A,_ and ProQ_V96D_. 5S RNA was detected as loading control. In A. and B. samples were obtained from cultures grown in LB to OD_600nm_ 2.0. **C**. Protein stability assays of pProQ WT and ProQ_V96D_. **D**. Protein stability assays of ProQ_F25S_, and ProQ_R80A_. **E**. Protein stability assay of ProQ_L71A._ In C, D and E cultures carrying the ProQ variants were grown in LB to OD_600nm_ 2.0. To stop translation, tetracycline was added to a concentration of 50 μg/ml. Samples for crude extracts were taken at time points 0, 15, 30, 60 and 120 minutes after tetracycline addition. ProQ-3×FLAG levels were determined by immunodetection. GroEL was detected as loading control. **F**. Quantification of panels C, D and E.

All the four mutations that caused reduced protein levels were located within the CDS of *proQ* and were therefore unlikely to affect protein synthesis. To fully rule out general expression effects, we probed the mRNAs of the four constructs with FinO domain mutations on a northern blot. The mutant plasmids (ProQ_F25S_, ProQ_L71A_, ProQ_R80A_ and ProQ_V96D_) produced even more *proQ* mRNA than did the plasmid expressing wild-type ProQ (Fig. 4B).

There were two possible explanations for the lower protein levels: impaired translation of the mutant mRNAs or decreased stability of the mutant proteins. Because the mutations were in the CDS, impaired translation was unlikely. Accordingly, we analyzed the rate of protein decay after stopping cellular translation with the antibiotic tetracycline as previously described (34). As shown in Figure 4C, the wild-type ProQ protein is very stable *in vivo*, displaying almost no decay over the course of 120 min into the tetracycline treatment. By contrast, all four tested ProQ mutant proteins (ProQ_F25S_, ProQ_L71A_, ProQ_R80A_ and ProQ_V96D_) display significantly reduced half-lives (Fig. 4C-E). Quantification of these western blot results revealed a dramatic decrease in protein half-life, from >120 min (WT ProQ) to ∼11 min for the ProQ_R80A_ and ProQ_V96D_ mutants, ∼5 min for the ProQ_F25S_ mutant, and a milder reduction to ∼60 min for ProQ_L71A_ (Fig. 4F). The reduced half-lives correlated well with the lower steady-state levels of these mutant proteins (see Fig. 4A). Our findings show that the mutations do not only impair RNA-binding, but also the intrinsic stability of the ProQ protein *in vivo*.

### Lon protease degrades ProQ mutant proteins

Protein turnover in enteric Gram-negative bacteria is primarily driven by members of the adenosine triphosphatase (ATPase) associated with cellular activities (AAA+) family. These proteases turn over functional proteins and also remove potentially toxic non-functional proteins (35–37). To determine which protease was responsible for clearing the non-functional ProQ proteins, we selected three major proteases from this family: ClpA, ClpX and Lon. Probing for the ProQ_F25S_, ProQ_R80A_ and ProQ_V96D_ mutant proteins in individual protease knockout strains of *Salmonella*, we observed full restoration to wild-type levels for all three mutant proteins in the Δ*lon* strain (Fig. 5A). By contrast, only minor changes were observed in the Δ*clpA* or Δ*clpX* strains (Fig. 5B-C).

**Figure 5.**
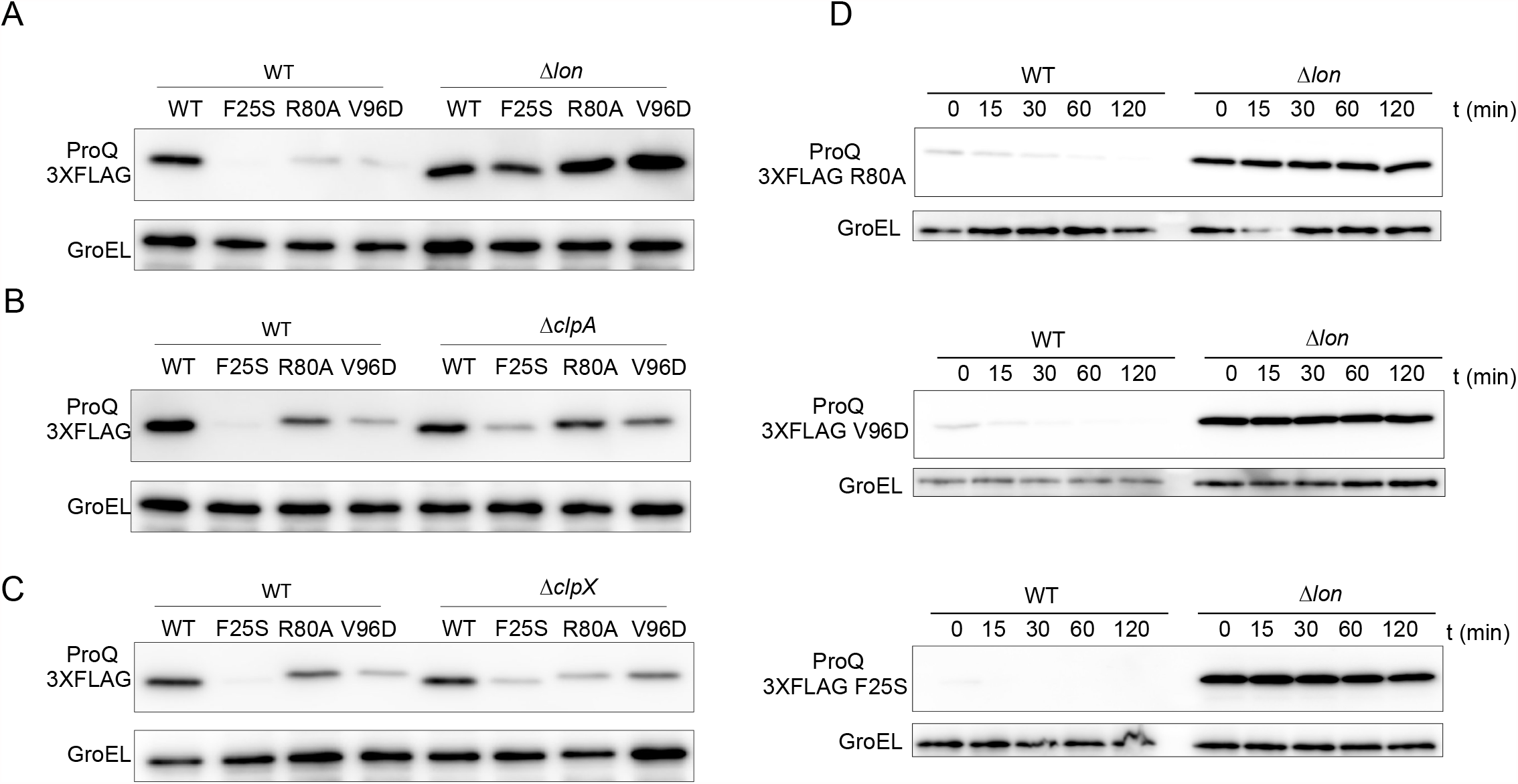
ProQ inability to interact with RNA leads to Lon-mediated proteolysis. Immunodetection of ProQ-3×FLAG variants pProQ WT, ProQ_V96D,_ ProQ_F25S_, and ProQ_R80A_ in **A**. WT and Δ*lon* (upper panel) **B**. WT and Δ*clpA* (middle panel) **C**. WT and Δ*clpX* (bottom panel). GroEL was detected as loading control. Absence of Lon but not ClpA or ClpX affects steady state levels of ProQ_V96D,_ ProQ_F25S_, and ProQ_R80A_ when compared to pProQ WT. **D**. Protein stability assays in WT and Δ*lon* of i) pProQ WT and ProQ_R80A_ (upper panel) ii) pProQ WT and ProQ_V96D_ (mid panel) and pProQ WT and ProQ_F25S_ (bottom panel). Cultures carrying the ProQ variants were grown in LB to OD_600nm_ 2.0. To stop translation, tetracycline was added to a concentration of 50 μg/ml. Samples for crude extracts were taken at time points 0, 15, 30, 60 and 120 minutes after tetracycline addition. ProQ-3×FLAG levels were determined by immunodetection. GroEL was detected as loading control.

The crucial involvement of Lon for removing non-functional ProQ proteins was corroborated by the results of *in vivo* protein stability experiment. Specifically, we observed a dramatic increase in protein half-life for the ProQ_F25S_, ProQ_R80A_ and ProQ_V96D_ in the Δ*lon* strain (Fig. 5D), essentially restoring it to the wild-type situation (t _1/2_ >120 min). These observations suggested a mechanism whereby ProQ variants unable to interact with RNA are actively targeted by the protease Lon and rapidly cleared from the cell.

### Evidence that ProQ mutant proteins are defective in RNA-binding

Our assumption that the selected FinO-domain mutants ProQ are defective in RNA binding was based on both, inference from an *E. coli* 3HB screen by others (26) and the strong reduction of reduced SibA RNA levels observed here (Fig. 3C). The caveat being that these mutant proteins failed to accumulate *in vivo* (Fig. 4A), which limited conclusions regarding intracellular activity. In this regard, the Δ*lon* strain provided a unique opportunity to prove their inability to bind target RNAs *in vivo*. Indeed, northern blot analysis showed that despite the increase in ProQ protein levels, SibA RNA levels remained low in the Δ*lon* strain, as compared to Lon-proficient *Salmonella* (Fig. 6A). In other words, these FinO-domain mutants could not stabilize the SibA RNA target even when present at wild-type concentration.

**Figure 6.**
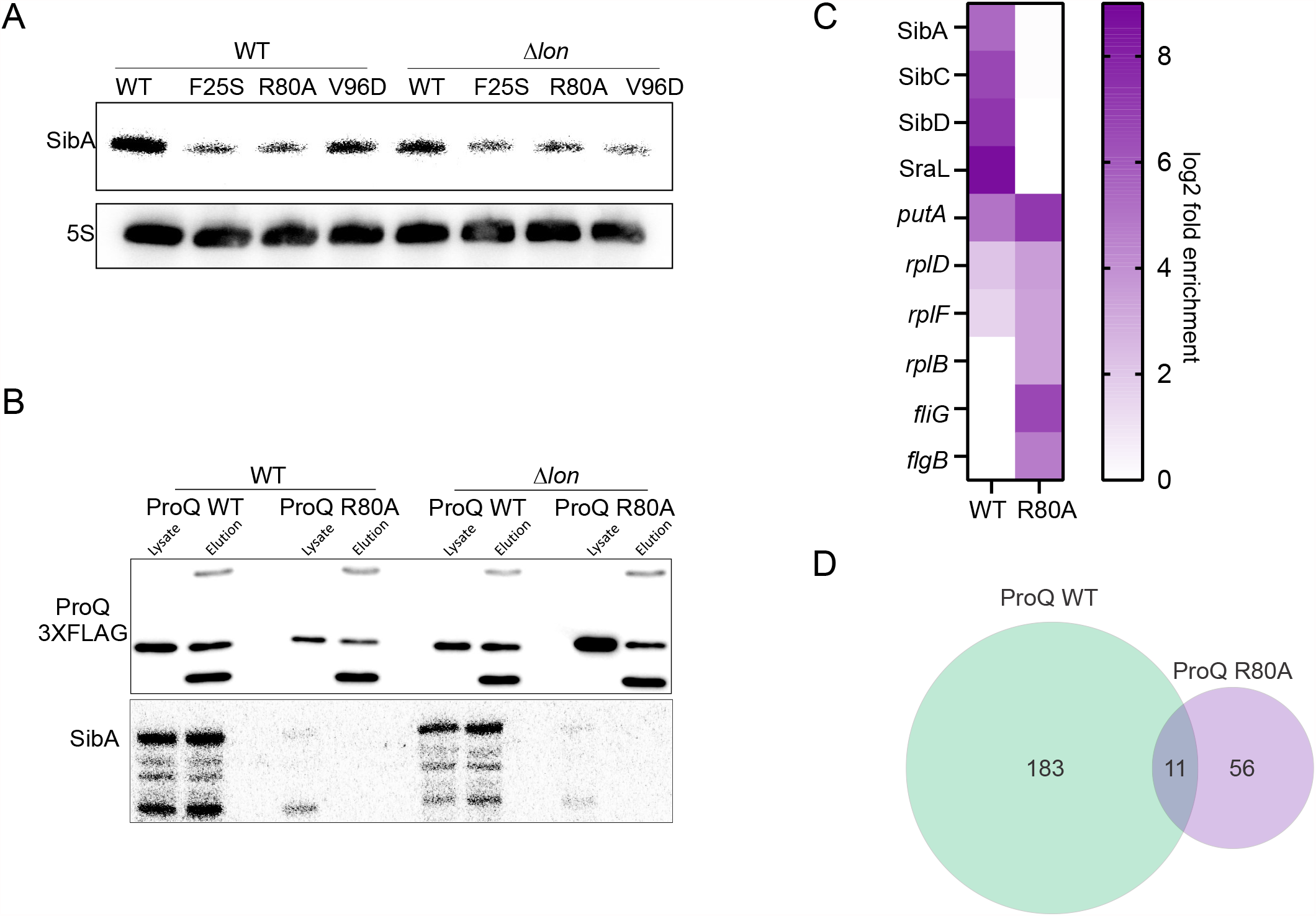
ProQ non-functional variants are unable to interact with RNA. **A**. Northern blot detection of SibA sRNA expression in Δ*proQ* background expressing pProQ variants: pProQ WT, ProQ_F25S_, ProQ_R80A_ and ProQ_V96D_ in WT (left) and Δ*lon* (right) deletion background. 5S RNA was detected as loading control. Samples were obtained from cultures grown in LB to OD_600nm_ 2.0. **B**. Immunoprecipitation of pProQ WT and ProQ_R80A_ in WT (left) and Δ*lon* (right) mutant background, lysate and elution extracts of protein (upper panel) and RNA (bottom panel) were prepared. Immunodetection (upper panel) of pProQ WT and ProQ_R80A_ protein variants immunoprecipitated. Northern blot detection (bottom panel) of co-immunoprecipitated SibA sRNA. **C**. Heatmap of highly enriched transcripts in RIP-seq of pProQ WT and ProQ_R80A_ **D**. Venn diagram of significantly enriched transcripts (log_2_fold ≥ 2.0; *p*-adj ≤ 0.05) in pProQ WT and ProQ_R80A_ in Δ*lon* background compared to VC strain.

In order to prove loss of RNA binding more directly, we selected the ProQ_R80A_ mutant for RNA co-immunoprecipitation (coIP) *in vivo*. As shown in Fig. 6B, coIP with the wild-type ProQ protein recovered with an α-FLAG antibody strongly enriched the SibA RNA, whereas no enrichment was observed with the mutant protein. Next, we globally analyzed the transcripts from the eluate fractions by RNA-seq. Precipitation with wild-type ProQ recovered the expected suite of dominant ProQ-associated sRNAs, e.g. SibA, SibD, SibC and SraL (Fig. 6C); in other words, the plasmid-expressed ProQ protein showed the same target profile as in previous coIP studies with ProQ expressed from the chromosome (9, 23). By contrast, ProQ_R80A_ had a more restricted target suite than wild-type ProQ, and enriched entirely different transcripts (Fig. 6D). In fact, ProQ_R80A_ associates nonspecifically with abundant mRNAs of ribosomal proteins, flagella components and even its own messenger (Fig. 6C-D, S3A). A clustering analysis revealed that the RNA profile of ProQ_R80A_ overall is similar to the coIP with the empty vector control (Fig. S3B). Thus, the R80A substitution renders ProQ unable to bind ProQ dependent RNAs, a deficiency that is associated with rapid degradation by the Lon protease.

### Hfq is also targeted by Lon

ProQ is one of two major RBPs to facilitate sRNA activity in *Salmonella*, the other being the hexameric Sm-like protein, Hfq (15, 20). Assembled Hfq hexamers interact with RNA through three different RNA-binding surfaces: their proximal and their distal face, and the rim. Residue substitutions rendering *E. coli* Hfq unable to interact with a subset of RNA species have been reported for each of these three surface regions (38–42). Intriguingly, some of these mutant proteins, e.g., Hfq_D40A_ and Hfq_Y55A_, also exhibited reduced protein levels in *E. coli* (43). Here, when we generated the corresponding mutations in Hfq from *Salmonella* expressed from a plasmid, we also observed a reduction in protein expression of Hfq_D40A_ and Hfq_Y55A_ as compared to wild-type Hfq (Fig. 7A). As a control, we generated the distal face mutant Hfq_Y25D_ which accumulated to wild-type Hfq levels (Fig. 7A), as previously reported in *E. coli* (43).

**Figure 7.**
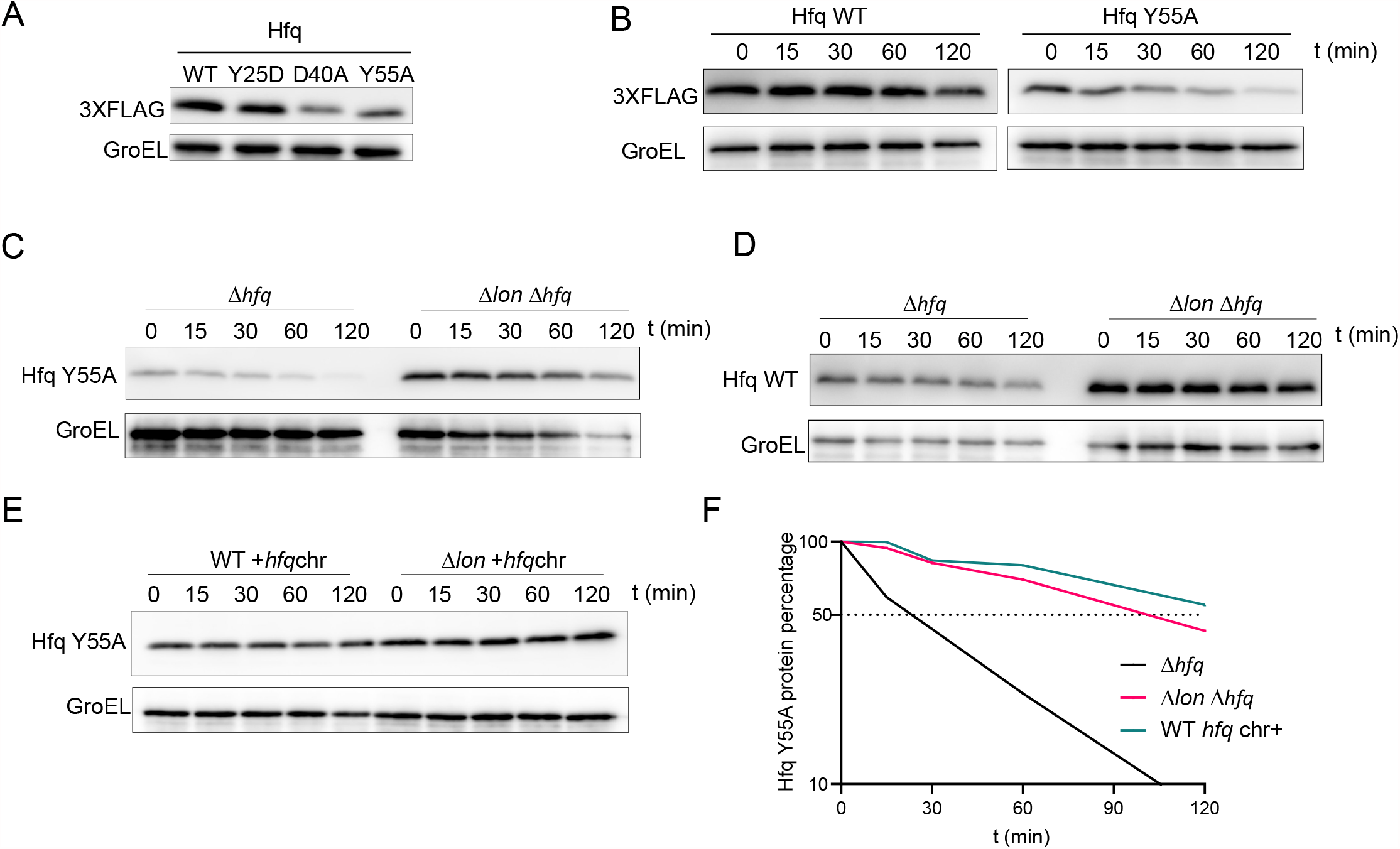
Lon targets the RBP Hfq. **A**. Immunodetection of Hfq-3×FLAG variants pHfq_WT_, Hfq_Y25D,_ Hfq_D40A_ and Hfq_Y55A_. GroEL was detected as loading control. **B**. Protein stability assays of pHfq_WT_ and Hfq_Y55A_ in a Δ*hfq* background **C**. Protein stability assays in Δ*hfq* and Δ*hfq*Δ*lon* backgrounds of Hfq_Y55A_. **D**. Protein stability assays in Δ*hfq* and Δ*hfq*Δ*lon* backgrounds of Hfq_WT_. **E**. Protein stability assays in WT and Δ*lon* backgrounds of Hfq_Y55A_. **F**. Quantification of panels C and E. In B, C, D and E cultures carrying the Hfq variants were grown in LB to OD_600nm_ 2.0. To stop translation, tetracycline was added to a concentration of 50 μg/ml. Samples for crude extracts were taken at time points 0, 15, 30, 60 and 120 minutes after tetracycline addition. Hfq-3×FLAG levels were determined by immunodetection. GroEL was detected as loading control.

To test a potential role of Lon in quality control of RNA-binding deficient Hfq variants, we selected the Hfq_Y55A_ mutant for stability analysis. Similar to ProQ, wild-type Hfq displayed a half-life of ≥120 min in our tetracycline treatment assay, whereas the intracellular half-life of the Hfq_Y55A_ protein was much shorter (Fig. 7B). Moreover, we observed for the Hfq_Y55A_ protein the same clear rescue of intracellular stability in the Δ*lon* mutant (Fig, 7C) as we had with the RNA-binding deficient ProQ mutants above. Wild-type Hfq was stable over the course of 120 min, and an additional increase in Hfq protein levels was observed in the Δ*lon* mutant (Fig. 7D).

The Hfq_Y55A_ variant is impaired in hexamer formation, which requires interaction with RNA (39, 43, 44). We hypothesized that the reduced intracellular stability of Hfq_Y55A_ was due to the inability of this mutant protein to form the active hexamer. Interestingly, it has been shown that wild-type Hfq monomers can form heterohexamers with Hfq mutant variants, and thereby rescue interaction with RNA in *E. coli* (39, 44). Mimicking this approach, we expressed Hfq_Y55A_ in a *Salmonella hfq*+ background, expecting that heterohexamers would form and lead to stabilization of the mutant protein. We used the 3xFLAG tag on the mutant protein for selective detection by western blot. We discovered that the half-life of the mutant Hfq_Y55A_ protein dramatically increased to ≥120 min, making it indistinguishable from wild-type Hfq, regardless of the presence or absence of Lon (Fig. 7E-F).

## DISCUSSION

The global and specialized roles played by RBPs of the ProQ/FinO family reflect posttranscriptional control mechanisms that have been discovered recently in a number of bacteria, including *E. coli, Legionella pneumophila, Neisseria meningitidis*, and *S. enterica* (9, 11, 12, 14, 15, 23). Although it was originally proposed that ProQ only functioned as a positive regulator of proline transport, it was subsequently discovered that this RNA chaperone targeted a large number of cellular transcripts (45). It has been inferred from a combination of protein conservation, biochemical and genetic studies that the key regulatory function of ProQ is associated with the FinO domain and not the variable C-terminal region. However, this assumption needed to be tested experimentally. Our present study pioneers a saturating mutagenesis screen over the entire length of ProQ under physiological conditions. We selected mutations that inactivate ProQ, based on the ability of the RBP to repress microaerobic growth of *Salmonella* on succinate.

One unexpected finding from our screen was that certain residues in the variable C-terminus are required for ProQ function. We identified two validated C-terminal mutations, a change of glutamine 185 to valine, and leucine 188 to proline, that lie in a region which has been predicted to be involved in RNA binding by a protein structure modeling approach (22). However, these mutations only had a modest influence on SibA RNA, a sensitive reporter of ProQ activity (Fig. 3C), resembling the impact of frameshift mutations in the C-terminal domain of the related RocC protein in *Legionella* which also impaired regulation of RocC targets without affecting RNA-protein interactions (11).

Until now, the evidence for an independent role for the ProQ CTD was circumstantial as some RNA targets of *E. coli* ProQ are bound more efficiently by the full-length protein than by the NTD alone (27). To improve our understanding of the C-terminal domain, we suggest that other loss-of-function approaches should be used in future. An obvious selectable phenotype would be resistance to 3,4-dehydroproline, which was the first indication of the physiological role of ProQ (45, 46). If the same C-terminal residues played a functional role that involved a different aspect of ProQ activity, it would strengthen the idea that the C-terminus constitutes an important domain in its own right.

As well as revealing an unexpected function for the CTD, our scanning mutagenesis over the full-length protein emphasizes the importance of the N-terminal FinO domain for ProQ function *in vivo*. The screen predicts 27 residue substitutions within this domain that lead to a non-functional ProQ variant, consistent with the 25 residue substitutions reported in the recently published three-hybrid (3HB) based screen, which used just the NTD of *E. coli* ProQ and *cspE* or SibB RNAs as a molecular bait (26). While most of the described amino acid substitutions had overlapping functions for both *cspE* and SibB, some specificity was observed (e.g., ProQ_K35E_ and ProQ_R20P_ only influenced SibB binding).

One possible limitation of the *E. coli* 3HB screen was that the NTD was expressed as a fusion protein with the alpha subunit of RNA polymerase, and was not investigated in the context of the entire ProQ protein. Overall, a strong overlap between the mutations we identified in the *Salmonella* ProQ NTD with the mutations from the *E. coli* 3HB screen was observed. Key differences include *E. coli* ProQ_K35E_, a mutation that was not enriched in the *Salmonella* library, and our *Salmonella* ProQ_V96D_ mutant, which was not detected in the *E. coli* screen (Fig. S4, Supplementary Table S2). These observations indicate subtle differences between the RNA-binding faces of the *E. coli* and *Salmonella* ProQ proteins, despite the high level of similarity between these proteins.

The AAA+ protease Lon plays a crucial role in enteric bacteria by degrading important proteins to optimize particular regulatory functions (36, 37, 47). However, Lon is not the major degradation enzyme of ProQ, as wild-type ProQ does not accumulate to higher levels in the absence of Lon (Fig. 5A). Importantly, we have discovered several amino acid substitutions in the FinO domain that both abrogate RNA binding and to the resulting mutant ProQ proteins being rapidly degraded by Lon. This finding is consistent with a quality control mechanism of ProQ function *in vivo* that actively targets ProQ molecules that fail to associate with RNA (Fig. 8).

**Figure 8.**
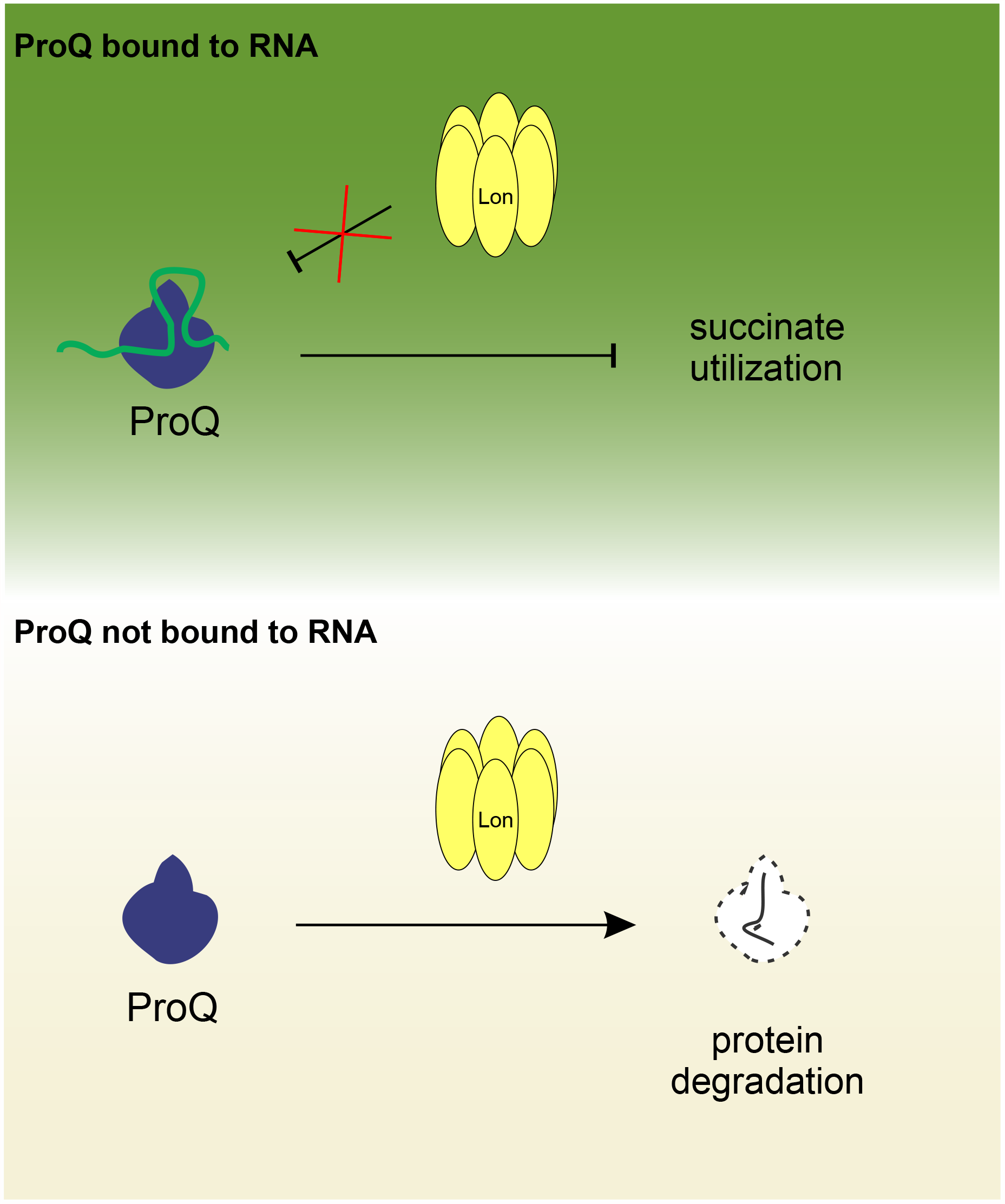
Role of Lon-mediated quality control of ProQ. When interacting with RNA, ProQ is stable as it escapes Lon-mediated degradation and carries out its *in vivo* function as suppressor of succinate utilization in *Salmonella* (upper panel). In the absence of RNA binding, however, ProQ is unstable as it is actively targeted by Lon (bottom panel).

To support our suggestion that Lon-mediated proteolysis is a general mechanism that prevents the accumulation of non-functional RBPs in bacteria, we show that Lon is also responsible for degrading an RNA-binding deficient variant of the sRNA chaperone Hfq. Furthermore, the Narberhaus laboratory recently showed that the Ffh protein component of the conserved signal recognition particle (SRP) is also targeted by Lon in the absence of binding with its major ligand, the 4.5S RNA (48). Collectively, these observations suggest that *in vitro* systems should be developed to prove directly whether the proposed Lon-mediated degradation of these RBPs is indeed dependent upon RNA-binding. While it is unclear at this stage whether *Salmonella* ProQ can be degraded by purified Lon protease, we note that the Hfq protein of *Pseudomonas aeruginosa* may be a useful experimental tool as it is easily degraded by Lon *in vitro* (49).

Lon is an important protease in bacteria that plays a central role in maintaining cellular protein homeostasis by removing misfolded proteins (50, 51) and controls the turnover of important regulatory proteins including H-NS, RcsA, SoxS and SulA (52–55). The accumulated knowledge about the many functions of Lon in the cell raises the possibility that Lon-mediated degradation of RBPs could be regulated by additional factors. For example, cleavage of H-NS by Lon only occurs in the absence of bound DNA (55). Furthermore, the bacterial toxin CcdA, which is part of a plasmid-borne toxin-antitoxin system, is only degraded by Lon when not bound to the antitoxin (37, 56–58). This would fit with our prediction that Lon degrades Hfq when the initial monomeric protein has not yet assembled into the active hexamer.

Our experimental evolution strategy was based upon the *Salmonella* succinate-related phenotype, which holds great promise for the design of additional genetic screens to search for novel factors that modulate the activity of Lon and ProQ, either alone or in combination.

## MATERIALS AND METHODS

### Bacterial strains and growth conditions

The bacterial strains and plasmids used in this study are listed in Supplementary Table S1. *Salmonella enterica* serovar Typhimurium SL1344 (59) and derivative strains were cultivated in Lysogeny broth (LB) (tryptone 10 g/l, yeast extract 5 g/l and sodium chloride 10 g/l) or mineral medium M9. Mineral M9 contained: i) 12.8 g/l Na_2_HPO_4_×7H_2_O, 3 g/l KH_2_PO_4_, 0.5 g/l NaCL, 1 g/l NH_4_Cl, 2mM MgSO_4_, 0.1 mM CaCl_2_, 0.0004% Histidine, 0.5 μg/ml Thiamine; ii) trace elements: 134 μM EDTA, 31 μM FeCl_3_×6H_2_O, 6.2 μM ZnCl_2_, 0.76 μM CuCl_2_×2H_2_O, 0.42 μM CoCl_2_×2H_2_O, 1.62 μM H_3_BO_3_, 81 nM MnCl_2_×4H_2_O; and iii) supplemented with 40 mM Na-succinate (Sigma-Aldrich) as carbon source. Bacterial cultures were inoculated to an OD_600 nm_ 0.01 and incubated at 37°C without aeration for microaerobic conditions as previously described (32, 60) or with 200 rpm shaking for aerobic conditions. For solid growth, *Salmonella* strains were streaked on LB or M9 Agar supplemented with 40mM Na-succinate as sole carbon source.

Growth curves were carried out in 96-well plates in a final volume of 200 µl. Plates were inoculated to an OD_600 nm_ 0.01 with cell suspensions of the strains of interest prewashed in 1×PBS. Plates were incubated at 37 °C without shaking for 24 hours in a Tecan Infinite M Plex plate reader. Growth rate was assessed by turbidity measurement (OD _600nm_) every 15 minutes.

### Genetic manipulations and site directed mutagenesis

*Salmonella enterica* deletion strains were generated by standard gene replacement as previously described (61). Generated PCR fragment is transformed in competent *Salmonella* as for deletion strains (61). When required, the antibiotic cassette was removed by the expression of Flp recombinase from pCP20 as previously described (62).

A *proQ* and *proQ*-3×FLAG variant was cloned in between AatII/XbaI in pZE12luc backbone, to be expressed under *proQ* native promoter with oligonucleotide pairs JVO-16806/JVO-8524 and JVO-16806/JVO-12604 respectively. The resulting expression plasmid pProQ-3×FLAG was used as template for generation of ProQ point mutation variants. The mutations were generated by site directed mutagenesis. Primers of 40 bp that contain the substitution of interest were used to amplify by PCR pProQ-3×FLAG. The following mutations were generated with the indicated oligonucleotides pairs: ProQ_F25S_ (JVO18952/JVO18953), ProQ_R80A_ (JVO18550/JVO18551), ProQ_L71A_ (JVO18552/JVO18553), ProQ_V96D_ (JVO18954/JVO18955), ProQ_Q118R_ (JVO18956/JVO16957), ProQ_K148E_ (JVO18960/JVO18961), ProQ_L188P_ (JVO18962/JVO18963), ProQ_V190E_ (JVO18964/JVO18965), ProQ_Q212R_ (JVO18966/JVO18967), ProQ_V227E_ (JVO18968/JVO18969), ProQ_G185V_ (JVO19480/JVO19481), ProQ_I204V_ (JVO19482/JVO19483), ProQ_E203G_ (JVO19484/JVO19485). The resulting reaction was DpnI-treated and transformed into chemo competent *E. coli* TopF. Mutant candidates were validated by Sanger sequencing.

Similarly, *hfq*-3×FLAG variant was cloned between XhoI/XbaI in pZE12luc backbone to be expressed under *hfq* native promoter with oligonucleotides pairs JVO19817/JVO19818. The resulting expression plasmid pHfq3×FLAG was used as a template to generate Hfq point mutations variants as described above for ProQ. Oligonucleotides used for strains construction, cloning and site directed mutagenesis are listed in Supplementary Table S1.

### Protein crude extracts and western blot

Bacterial cultures were grown to OD_600 nm_ 2.0. A volume of cells that represents 0.2 OD_600 nm_ were pelleted and resuspended in 200 μL of 1× Laemmli buffer. Samples were stored at –20°C. Protein extracts were subjected to SDS-PAGE separation and transferred to a PVDF filter membrane. The membrane was subjected to immunodetection of 3×FLAG tagged proteins by using as primary antibodies, monoclonal anti-FLAG M2 (Sigma-Aldrich #F1804) 1:2000 for both ProQ-3×FLAG and Hfq-3×FLAG detection. As loading control, GroEL was detected with anti-GroEL (Sigma-Aldrich, #G6532). As of secondary antibodies, anti-mouse (Thermo Fisher, cat# 31430) and anti-rabbit (Thermo Fisher, cat# 31460) conjugated to horseradish peroxidase were used for anti-FLAG and anti-GroEL respectively. For detection, ECL_TM_ Prime Western Blotting Detection Reagent (Cyvita) served as a substrate.

### In vivo protein stability assays

Bacterial cultures were grown to OD_600 nm_ 2.0. To stop translation, tetracycline was added to the culture to a concentration of 50 μg/ml as previously described (34). Time points were collected, prior to the addition of the antibiotic, t0, and after 15, 30, 60, 120 minutes. For each time point, 96 μL of culture were mixed with 24 μL of 5× Laemmli buffer. Samples were stored at –20°C. The levels of ProQ-3×FLAG WT, Hfq-3×FLAG WT and mutant variants were determined by immunodetection by western blot as described above.

### Total RNA isolation and northern blot

Bacterial cultures were grown to OD_600 nm_ 2.0, a volume of cells that represents 4 OD_600 nm_ was collected, and total RNA extracted by hot phenol method followed by a DNase I treatment. Samples of 10 μg of DNAse-treated total RNA were subjected to electrophoretic separation in Tris-Borate-EDTA (TBE) 6% acrylamide gels containing 8.3 M urea. RNAs were transferred to Hybond N+ (GE Healthcare) filters and transcripts of interest (SibA sRNA and *proQ* mRNA) were detected by hybridization with 5’ radiolabeled oligonucleotides as probes. For oligonucleotides labeling, in a 20 μL reaction, 10 pmol of the oligonucleotides used as probes were 5’-labeled with 10 μCi of _32_P-γ-ATP by PNK (T4 polynucleotide kinase, Thermo Fisher Scientific) for 1 h at 37°C. Labeled oligonucleotides were further purified via Microspin G-25 columns (GE Healthcare) to remove unincorporated _32_P-γ-ATP. Radioactive signal was imaged with the Typhoon FLA 7000 (GE Healthcare). Oligonucleotides used as probes are listed in Supplementary Table S1.

### Deep mutational scanning and DNA library preparation

Libraries of ProQ mutants were generated by using GeneMorph II EZClone Domain Mutagenesis Kit (Agilent). As template, three different overlapping libraries were generated, LIB1 (1-81 aa), LIB2 (73-155 aa) and LIB3 (155-227 aa). First a PCR template was generated for each library by amplifying overlapping regions of the *proQ* ORF with JVO16789/JVO16809, JVO16810/JVO16728 and JVO16811/JVO16790 for LIB1, LIB2 and LIB3 respectively, generating 240, 250 and 218 base pair PCR templates. For each library a PCR fragment was generated by error prone PCR following Agilent guidelines for low mutation rate (0-4.5 mutations/kB) (Fig. S2). The purified PCR products were subsequently used as megaprimers to generate the library of mutants in the plasmid pZE-ProQ-3×FLAG by following GeneMorph II EZClone Domain Mutagenesis Kit (Agilent) guidelines. For each library, around 5000 colonies were obtained upon transformation of DpnI-treated mutagenesis reaction into chemo competent *E. coli* TopF cells. All transformants were pooled and mutant plasmid library was extracted by plasmid Midi prep (Qiagen) and transformed by electroporation into Δ*proQ* cells.

Around 30,000 transformants/library were obtained and pooled in 10 ml 1×PBS. Cells were washed twice on fresh 1×PBS prior to inoculation on the selective (M9+succinate) and non-selective (LB) media. The turbidity of the transformants suspension was measured and 4 ODs were stored as ‘input’. Next, 2 ODs were inoculated in 50 ml of either M9+succinate or LB in a 250 ml flask and incubated statically (without aeration) for 22h at 37°C (Fig. 3C). From the resulting cultures, 4 ODs were collected and considered ‘output’. Plasmid content from ‘input’ and ‘output’ was extracted by Nucleospin Mini plasmid kit (Macherey-Nagel) and used as template for library preparation for deep sequencing as detailed below.

Library preparation was carried out by using 50 ng of plasmid as template from the ‘input’ and ‘output’. For each library LIB1 (1-81 aa), LIB2 (73-155 aa) and LIB3 (155-227 aa), same region subjected to mutagenesis was amplified by PCR with specific primers that incorporated Nextera adapters: i) JVO18138/JVO18139 for LIB1, ii) JVO18140/JVO18141 for LIB2 and iii) JVO18142/JVO18143 for LIB3. The distribution of the library and concentration of library DNA was determined by DNA bioanalyzer and a DNA Qubit measurement respectively. Amplified DNAs from different libraries were pooled and sequenced on an Illumina Mi-seq pair ended 300 bp at the Next generation sequencing core unit of the Helmholtz Centre for Infection Biology (HZI).

### Deep mutational scanning data analysis

Deep mutational scanning generated read counts were analyzed by applying Enrich2 statistical framework (63). Shortly, for each library, read counts were mapped to ProQ mutagenized region LIB1 (1-81 aa), LIB2 (73-155 aa) and LIB3 (155-227 aa) respectively. Enrichment factors were calculated between the ‘output’ and ‘input’ libraries in M9 succinate (one biological duplicate each) and in LB (28, 63). The reads were used as input for Enrich2 (63) to determine enrichment of point mutations. For this, Enrich2 settings were set to include reads with an average read quality above 20 and up to 50 mismatches to the ProQ sequence and unmatched paired reads were excluded. The enrichment scores and Z-scores (deviation from the overall mean) were calculated using the setting “log ratios (Enrich2)” after normalization to the detected wild type sequences.

### RNA-coimmunoprecipitation

RNA-coimmunoprecipitation was carried out as previously described. *Salmonella enterica* WT and derivative Δ*lon* strains carrying either a vector control (VC), pProQ-3×FLAG WT or pProQ-3×FLAG R80A variants were grown in LB to OD_600 nm_ 2.0. A volume of cells representing 50 ODs were pelleted and cells resuspended in 800 μL of lysis buffer (20 mM Tris pH8.0, 150 mM KCl, 1 mM MgCl_2_, 1 mM DTT) supplemented with 8 μL of DNase I. The cells were lysed through mechanical lysis by 0.1 mm glass beads at 30 Hz for 10 min in the Retsch MM200. The lysate was cleared by centrifugation. For protein input control, a volume equivalent to 0.5 OD_600 nm_ was diluted to 90 μL with 1× Laemmli buffer (lysate protein sample) and stored at -20°C. For RNA input control, a volume equivalent to 5 OD_600 nm_ was saved for RNA extraction with TRIzol (lysate RNA sample). The remaining lysate was then incubated with 25 μL (1 μL/2 OD_600 nm_ of cells) of monoclonal anti-FLAG M2 (Sigma #F1804) at 4°C with rotation for 30 min. The sample was then added to 75 μL of pre-washed Protein A sepharose beads (Sigma-Aldrich) and further incubated with rotation at 4 °C for an additional 30 min. The beads were washed with 500 μL of lysis buffer for five times and finally resuspended in 532 μL of lysis buffer for elution of ProQ 3xFLAG variants and co-immunoprecipitated RNA.

For protein elution control, to a volume of 32 μl, 8 μL of 5× Laemmli buffer was added and stored at -20°C (elution protein sample). For RNA elution, co-immunoprecipitated RNA was extracted by phenol:chloroform:isoamyl alcohol (P:C:I) (25:24:1, pH 4.5, Roth) extraction. The purified RNA samples were treated by DNase I and subsequently purified by P:C:I extraction. Samples were resuspended in H_2_O to a final concentration of 1 OD/μL for elution samples and 0.1 OD/μL for lysate, flowthrough and wash. Resulting RNA samples were subjected to visualization via northern blot or quantification via deep sequencing. Protein extracts were also analyzed by western blot.

### cDNA library preparation and RIP-seq data analysis

Library preparation from co-immunoprecipitated RNA samples were carried out with NEBNext Multiplex Small RNA Library Prep Set (New England Biolabs) that allow sequencing in Illumina platforms. The library preparation was carried out in a thermocycler and manufacturer’s guidelines were followed with minor modifications as previously described (23). The distribution of the library and concentration of library DNA was determined by DNA bioanalyzer and a DNA Qubit measurement respectively. Amplified cDNAs from different libraries were pooled and sequenced on an Illumina NextSeq 500 platform at the Core Unit SysMed at the University of Würzburg. Mapping and quantification was performed as previously described (23).The enrichment of RNA was calculated by comparing co-enriched RNA with ProQ-3×FLAG WT and ProQ-3×FLAG R80A protein variants to the vector control (VC) strain. This experiment was performed in biological duplicates (Supplementary Table S3).

## DATA AVAILABILITY

The sequencing data have been deposited in NCBI Gene Expression Omnibus (64) and are accessible through GEO Series accession number GSE174509. Reviewers can access the dataset with the secure token ‘evebqmiubxkhvkd’.

## ACKNOWLEDGMENTS

We thank Jens Hör, Gianluca Matera and Susan Gottesman for fruitful discussions. We are grateful to Svetlana Durica-Mitic and Gianluca Matera for critical reading of the manuscript. We thank Eduardo Groisman and Josep Casadesus for providing strains. We appreciated advice on Lon-mediated degradation from Franz Narberhaus and Simon Brückner. We thank the Vogel Stiftung Dr. Eckernkamp for supporting F.P. with a Dr. Eckernkamp Fellowship.

## FUNDING

This work was supported by a DFG Leibniz Award to J.V. (Vo875-18), and, in part, by a Wellcome Trust Senior Investigator award to J.C.D.H. [Grant number 106914/Z/15/Z]. For the purpose of open access, the author has applied a CC BY public copyright licence to any Author Accepted Manuscript version arising from this submission.

## Notes

### Competing Interest Statement

The authors have declared no competing interest.

