## Supplementary material for "Scanning mutagenesis of RNA-binding protein ProQ reveals a quality control role for the Lon protease"

#### Supplementary figures legends

**Figure S1. ProQ represses succinate utilization through the TCA cycle.** Growth of *Salmonella* WT, Δ*proQ*, Δ*sdhA* and Δ*sdhA*Δ*proQ* in a 96-well plate Tecan. Strains were inoculated to an $OD_{600\ nm}$ 0.01 and grown in a 96-well plate in M9 40mM succinate at 37°C for 24 hours (h) without shaking. Growth rate was assessed by turbidity measurement ($OD_{600nm}$) every 15 minutes.

**Figure S2. ProQ mutants library preparation.** Schematic representation of ProQ point mutations variants library preparation. Adapted from GeneMorph II EZClone Domain Mutagenesis Kit (Agilent) guidelines. Generated plasmids containing the point mutations were transformed in electrocompetent Δ*proQ* cells to generate the INPUT libraries.

**Figure S3. RIP-seq of pProQ$_{WT}$ and pProQ$_{R80A}$. A.** Volcano plot of RNA transcripts enriched by RIP-seq of ProQ$_{WT}$-3×FLAG (left panel) and ProQ$_{R80A}$-3×FLAG (right panel). **B.** PCA plot distribution of duplicate samples of ProQ$_{WT}$-3×FLAG and ProQ$_{R80A}$-3×FLAG RIP-seq samples shown in panel A and B.

**Figure S4. ProQ homologues alignment.** The protein sequence of i) ProQ from *Salmonella enterica* SL1344 (1) ii) RocC from *Legionella pneumophila* (2) and iii) ProQ from *Escherichia coli* K-12 (3) were aligned in Clustal Omega as a multiple sequence alignment and visualized in Jalview. Residues substitutions identified to impair ProQ/RocC function are highlighted in red. Highly enriched ProQ mutant variants identified in our screen are indicated in orange.

**Supplementary Table S1** Bacterial strains, plasmids and oligonucleotides.

**Supplementary Table S2** Deep mutation scanning of ProQ libraries.

**Supplementary Table S3** RIP-seq of $pProQ_{WT}$ and $pProQ_{R80A.}$

**Supplementary Table S1.** Strains, plasmids and oligonucleotides used in this study

| Strains | Genotype | Source |
|---|---|---|
| JVS-1574 | SL1344 WT | (4) |
| JVS-10317 | SL1344 Δ*proQ:frt* | (1) |
| YMS-71 | SL1344 WT pJV300 | This study |
| YMS-83 | SL1344 Δ*proQ:frt* pJV300 | This study |
| YMS-153 | SL1344 Δ*proQ:frt* pZE12-ProQ (*aatII/xbaI*) | This study |
| SV5608 | SL1344 *his+ sdhA*::cat | (5) |
| YMS-115 | SL1344 *sdhA*::cat | This study |
| YMS-116 | SL1344 Δ*proQ:frt sdhA*::cat | This study |
| YMS-156 | SL1344 Δ*proQ:frt* pZE12-ProQ-3xFLAG | This study |
| YMS-311 | SL1344 Δ*proQ:frt* pZE12-ProQ-3xFLAG F25S | This study |
| YMS-286 | SL1344 Δ*proQ:frt* pZE12-ProQ-3xFLAG L71A | This study |
| YMS-285 | SL1344 Δ*proQ:frt* pZE12-ProQ-3xFLAG R80A | This study |
| YMS-312 | SL1344 Δ*proQ:frt* pZE12-ProQ-3xFLAG V96D | This study |
| YMS-318 | SL1344 Δ*proQ:frt* pZE12-ProQ-3xFLAG Q118R | This study |
| YMS-319 | SL1344 Δ*proQ:frt* pZE12-ProQ-3xFLAG K148E | This study |
| YMS-367 | SL1344 Δ*proQ:frt* pZE12-ProQ-3xFLAG G185V | This study |
| YMS-368 | SL1344 Δ*proQ:frt* pZE12-ProQ-3xFLAG L188P | This study |
| YMS-313 | SL1344 Δ*proQ:frt* pZE12-ProQ-3xFLAG V190E | This study |
| YMS-369 | SL1344 Δ*proQ:frt* pZE12-ProQ-3xFLAG E203G | This study |
| YMS-370 | SL1344 Δ*proQ:frt* pZE12-ProQ-3xFLAG I204E | This study |
| YMS-320 | SL1344 Δ*proQ:frt* pZE12-ProQ-3xFLAG Q212R | This study |
| YMS-314 | SL1344 Δ*proQ:frt* pZE12-ProQ-3xFLAG V227E | This study |
| EG18499 | *clpX*::cat | (6) |
| EG16039 | *lon*::cat | (6) |
| JY199 | *clpA*::cat | (7) |
| YMS-329 | SL1344 Δ*proQ:frt lon*::cat pZE12-ProQ-3xFLAG | This study |
| YMS-330 | SL1344 Δ*proQ:frt lon*::cat pZE12-ProQ-3xFLAG V96D | This study |
| YMS-331 | SL1344 Δ*proQ:frt lon*::cat pZE12-ProQ-3xFLAG R80A | This study |
| YMS-332 | SL1344 Δ*proQ:frt lon*::cat pZE12-ProQ-3xFLAG F25S | This study |
| YMS-333 | SL1344 Δ*proQ:frt clpA*::cat pZE12-ProQ-3xFLAG | This study |
| YMS-334 | SL1344 Δ*proQ:frt clpA*::cat pZE12-ProQ-3xFLAG V96D | This study |
| YMS-335 | SL1344 Δ*proQ:frt clpA*::cat pZE12-ProQ-3xFLAG R80A | This study |
| YMS-336 | SL1344 Δ*proQ:frt clpA*::cat pZE12-ProQ-3xFLAG F25S | This study |
| YMS-337 | SL1344 Δ*proQ:frt clpX*::cat pZE12-ProQ-3xFLAG | This study |
| YMS-338 | SL1344 Δ*proQ:frt clpX*::cat pZE12-ProQ-3xFLAG V96D | This study |
| YMS-339 | SL1344 Δ*proQ:frt clpX*::cat pZE12-ProQ-3xFLAG R80A | This study |
| YMS-340 | SL1344 Δ*proQ:frt clpX*::cat pZE12-ProQ-3xFLAG F25S | This study |
| YMS-377 | SL1344 Δ*hfq* pZE12-Hfq-3XFLAG | This study |
| YMS-387 | SL1344 Δ*hfq* pZE12-Hfq3XFLAG Y25D | This study |

| | | |
|---|---|---|
| YMS-388 | SL1344 Δ*hfq* pZE12-Hfq3XFLAG D40A | This study |
| YMS-389 | SL1344 Δ*hfq* pZE12-Hfq3XFLAG Y55A | This study |
| YMS-394 | SL1344 Δ*hfq*::cat Δlon pZE12-Hfq 3XFLAG | This study |
| YMS-399 | SL1344 Δ*hfq*::cat Δlon pZE12-Hfq 3XFLAG Y55A | This study |
| YMS-404 | SL1344 pZE12-Hfq 3XFLAG Y55A | This study |
| YMS-408 | SL1344 Δ*lon*::cat pZE12-Hfq 3XFLAG Y55A | This study |

| **Plasmids** | **Description** | **Source** |
|---|---|---|
| pKD46 | $Amp^R$ | (8) |
| pSUB11 | $Km^R$ | (9) |
| pJV300 | $Amp^R$ | (10) |
| pZE12-ProQ (*aatII*/*xbaI*) | Cloned in pZE12 backbone between AatII and XbaI. *proQ* is expressed under its native promoter. JVO16806/JVO8524. $Amp^R$ | This study |
| pZE12-ProQ-3xFLAG | Cloned in pZE12 backbone between AatII and XbaI. *proQ* is expressed under its native promoter as a 3XFLAG tagged variant. JVO-16806/JVO-12604. $Amp^R$ | This study |
| pZE12-ProQ-3xFLAG F25S | $Amp^R$ | This study |
| pZE12-ProQ-3xFLAG L71A | $Amp^R$ | This study |
| pZE12-ProQ-3xFLAG R80A | $Amp^R$ | This study |
| pZE12-ProQ-3xFLAG V96D | $Amp^R$ | This study |
| pZE12-ProQ-3xFLAG Q118R | $Amp^R$ | This study |
| pZE12-ProQ-3xFLAG K148E | $Amp^R$ | This study |
| pZE12-ProQ-3xFLAG G185V | $Amp^R$ | This study |
| pZE12-ProQ-3xFLAG L188P | $Amp^R$ | This study |
| pZE12-ProQ-3xFLAG V190E | $Amp^R$ | This study |
| pZE12-ProQ-3xFLAG E203G | $Amp^R$ | This study |
| pZE12-ProQ-3xFLAG I204E | $Amp^R$ | This study |
| pZE12-ProQ-3xFLAG Q212R | $Amp^R$ | This study |
| pZE12-ProQ-3xFLAG V227E | $Amp^R$ | This study |
| pZE12-Hfq-3xFLAG | Cloned in pZE12 backbone between XhoI and XbaI. *hfq* is expressed under its native promoter. JVO19817/JVO19818. $Amp^R$ | This study |
| pZE12-Hfq-3xFLAG Y55A | $Amp^R$ | This study |

| Oligonucleotides | Sequence |
|---|---|
| JVO-8524 | AGGCGTCTCTAGAAAAAAAAGTGTTCATGCCAGGCC |
| JVO-12604 | AGGCGTCTCTAGATTAATCATGATCCTTGTAGTCGATGTCATGATCTTTATAATCACCGT CATGGTCTTTGTAGTCGAACACCAGGTGTTCTGCGC |
| JVO-16806 | AGTTGTCGACGTCTACCGAAGAAGATGAACACGGCC |
| JVO-16966 | CATTCACGGCGAAGGTTATCGCTTCTGCGGCGACCTGCAGGATGACTACAAAGACCAT GACGG |
| JVO-16967 | TAACTTACCGGCTGTTTTTACAGTTTGGCGCCTGGGCCGAAGGTCCATATGAATATCCT CCTTAG |
| JVO-16789 | GTTGTAATCAGGAAATTTCATG |
| JVO-16809 | GCACGGATTGCCATCAAGGTC |
| JVO-16810 | CAAGCTGGCGTTACCTGTAC |
| JVO-16728 | CTTTCGTCGTCGGTTTATCAGCGC |
| JVO-16811 | GAAGGCGCTGAACGTAAACCTC |
| JVO-16790 | CAGGCCTGGCCTCCGTTTCA |
| JVO-18138 | TCGTCGGCAGCGTCAGATGTGTATAAGAGACAGGTTGTAATCAGGAAATTTCATG |
| JVO-18139 | GTCTCGTGGGCTCGGAGATGTGTATAAGAGACAGGCACGGATTGCCATCAAGGTC |
| JVO-18140 | TCGTCGGCAGCGTCAGATGTGTATAAGAGACAGCAAGCTGGCGTTACCTGTAC |
| JVO-18141 | GTCTCGTGGGCTCGGAGATGTGTATAAGAGACAGCTTTCGTCGTCGGTTTATCAGCGC |
| JVO-18142 | TCGTCGGCAGCGTCAGATGTGTATAAGAGACAGGAAGGCGCTGAACGTAAACCTC |
| JVO-18143 | GTCTCGTGGGCTCGGAGATGTGTATAAGAGACAGCAGGCCTGGCCTCCGTTTCA |
| JVO-18550 | GTTAAGCCGGGCGCAACGGCGGTCGACCTTGATGGCAATC |
| JVO-18551 | GATTGCCATCAAGGTCGACCGCCGTTGCGCCCGGCTTAAC |
| JVO-18552 | CTTTATACTTCAAGCTGGCGTTACGCGTACGGCGTTAAGCC |
| JVO-18553 | GGCTTAACGCCGTACGCGTAACGCCAGCTTGAAGTATAAAG |
| JVO-18952 | CCGAGCGTTTTCCTCACTGTTCTAGTGCGGAAGGCGAAGCT |
| JVO-18953 | AGCTTCGCCTTCCGCACTAGAACAGTGAGGAAAACGCTCGG |
| JVO-18954 | GAGCTGGAAGAACAGCATGACGAACATGCGCGTAAGCAGC |
| JVO-18955 | GCTGCTTACGCGCATGTTCGTCATGCTGTTCTTCCAGCTC |
| JVO-18956 | GCGCAGCGCGCAGAGCAGCGTGCGAAAAAACGCGAAGCTG |
| JVO-18957 | CAGCTTCGCGTTTTTTCGCACGCTGCTCTGCGCGCTGCGC |
| JVO-18960 | CTCGCCCGGTAGCGCGTCGTGAAGAAGGCGCTGAACGTAAAC |
| JVO-18961 | GTTTACGTTCAGCGCCTTCTTCACGACGCGCTACCGGGCGAG |
| JVO-18962 | GTATTGACCGTAGGGCAGTCCCCGAAGGTGAAAGCGGGTAATA |
| JVO-18963 | TATTACCCGCTTTCACCTTCGGGGACTGCCCTACGGTCAATAC |
| JVO-18964 | GTAGGGCAGTCCCTCAAGGAAAAAGCGGGTAATAATGCGA |
| JVO-18965 | TCGCATTATTACCCGCTTTTTCCTTGAGGGACTGCCCTAC |
| JVO-18966 | CAAAGATGGTGTCCGTGTACGCCTGAATTCGGGTATGTCT |
| JVO-18967 | AGACATACCCGAATTCAGGCGTACACGGACACCATCTTTG |
| JVO-18968 | GTACGCGCAGAACACCTGGAGTTCGACTACAAAGACCATG |
| JVO-18969 | CATGGTCTTTGTAGTCGAACTCCAGGTGTTCTGCGCGTAC |
| JVO-19480 | TATTTCAGTATTGACCGTAGTGCAGTCCCTCAAGGTGAAAG |
| JVO-19481 | CTTTCACCTTGAGGGACTGCACTACGGTCAATACTGAAATA |
| JVO-19482 | GGATGCCACCGTATTAGAAGTGACCAAAGATGGTGTCCGTG |

| | |
|---|---|
| JVO-19483 | CACGGACACCATCTTTGGTCACTTCTAATACGGTGGCATCC |
| JVO-19484 | GATGGATGCCACCGTATTAGGCATCACCAAAGATGGTGTCCG |
| JVO-19485 | CGGACACCATCTTTGGTGATGCCTAATACGGTGGCATCCATC |
| JVO-19817 | GTTCTCGAGGCTTGACAGTGAGAATCCCGATC |
| | AGGCGTCTCTAGATTAATCATGATCCTTGTAGTCGATGTCATGATCTTTATAATCACCGT |
| JVO-19818 | CATGGTCTTTGTAGTCTTCAGTCTCTTCGCTGTCCTGT |

### Figure S1

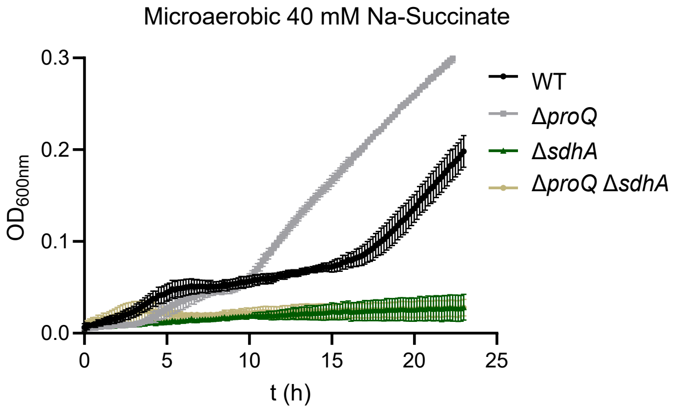
Microaerobic 40 mM Na-Succinate
0.3
0.2
0.1
0.0
OD600nm
0
5
10
15
20
25
t (h)
WT
ΔproQ
ΔsdhA
ΔproQ ΔsdhA

### Figure S2

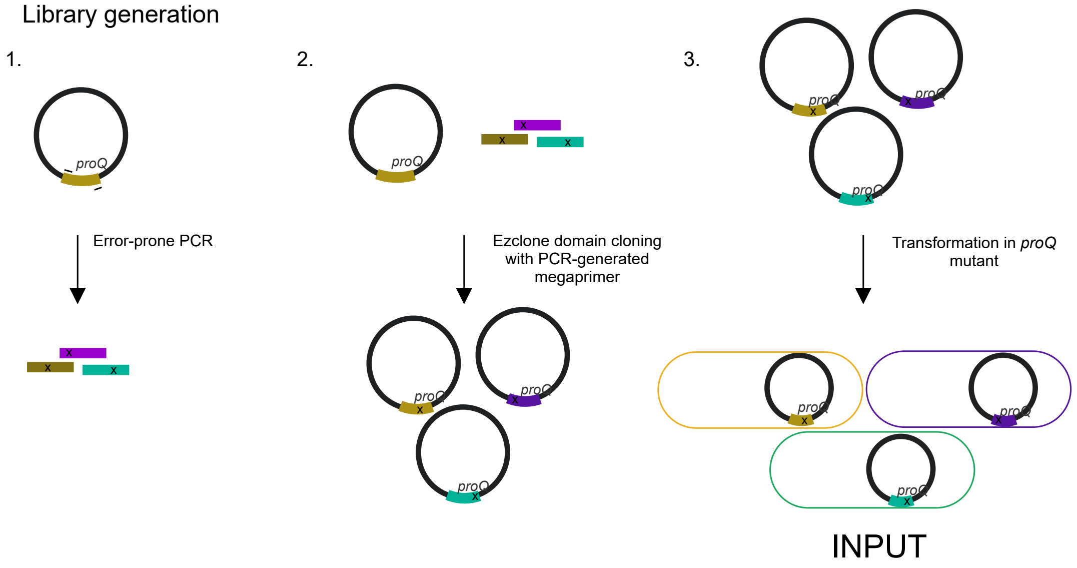

Library generation
1.
proQ
Error-prone PCR
x
x
x
2.
proQ
x
x
x
Ezclone domain cloning
with PCR-generated
megaprimer
proQ
x
proQ
x
proQ
x
3.
proQ
x
proQ
x
proQ
x
Transformation in *proQ*
mutant
proQ
x
proQ
x
proQ
x
INPUT

Figure S3

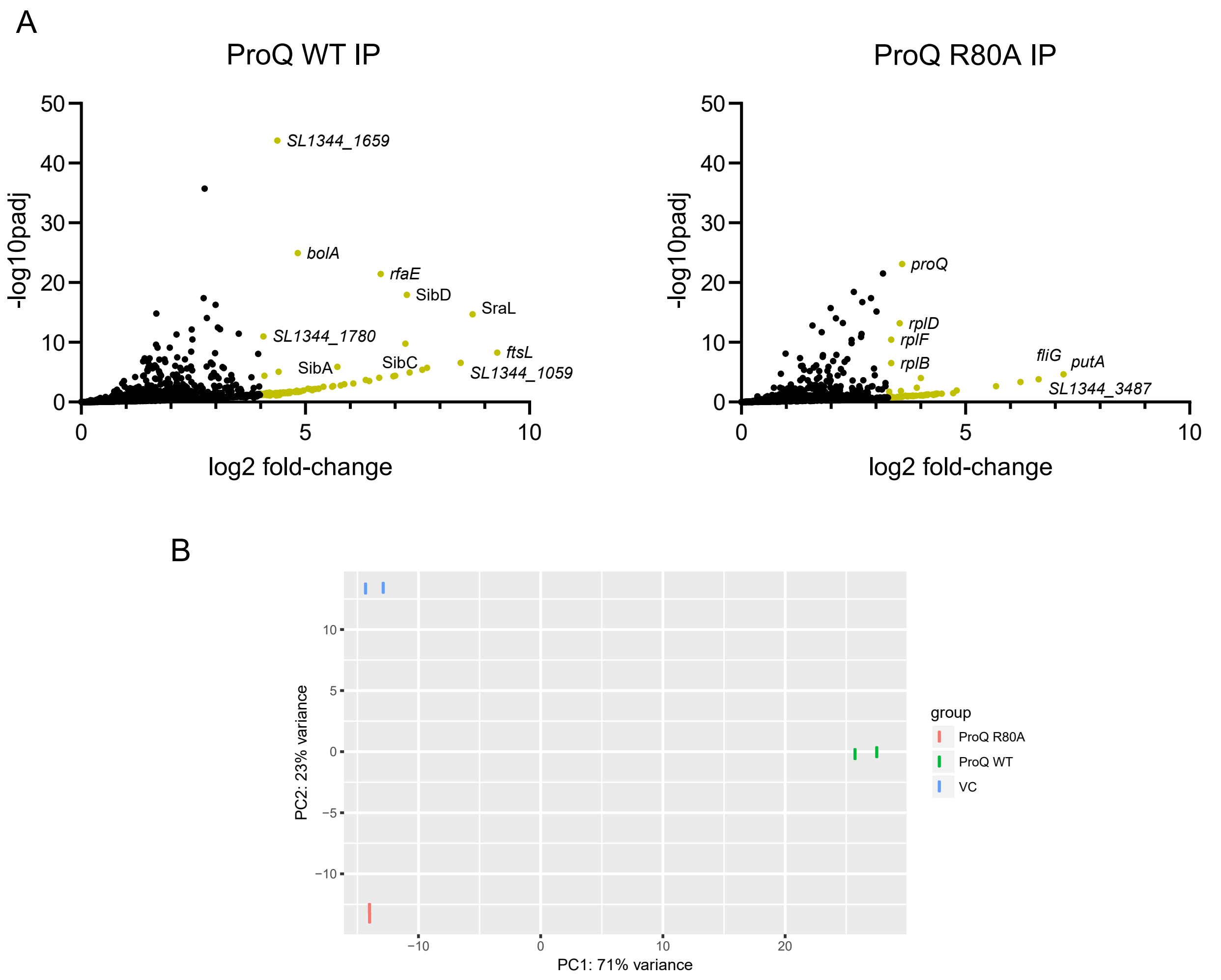

A
ProQ WT IP
-log10padj
log2 fold-change
SL1344_1659
bolA
rfaE
SibD
SraL
SL1344_1780
ftsL
SibA
SibC
SL1344_1059
ProQ R80A IP
proQ
rplD
rplF
rplB
fliG
putA
SL1344_3487
B
PC2: 23% variance
PC1: 71% variance
group
ProQ R80A
ProQ WT
VC

Figure S4

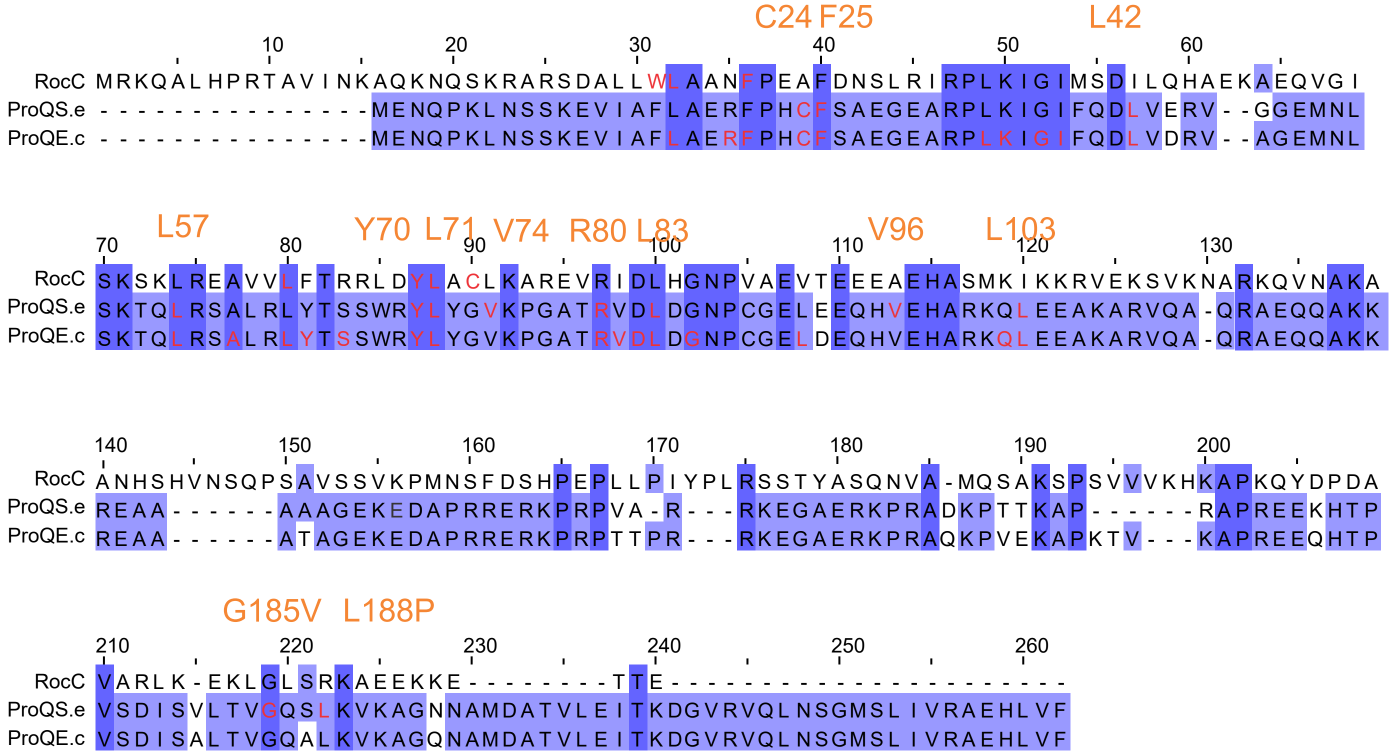

C24 F25
L42
RocC
ProQS.e
ProQE.c
L57
Y70 L71 V74 R80 L83
V96
L103
G185V L188P
